# First hematological and biochemical data in a rehabilitated giant pangolin (*Smutsia gigantea)* from southern Cameroon

**DOI:** 10.64898/2026.04.29.721582

**Authors:** Marine Montblanc, Jessica Harvey-Carroll, Julie Vanassche, Mark Donaldson, Ellen Connelly, Lisa Hywood

## Abstract

Giant pangolin (*Smutsia gigantea*) is one of the least studied pangolin species worldwide, with no published hematological and biochemical data available. We report the first blood parameters from a rehabilitated adult male from Campo Ma’an National Park (southern Cameroon). Hematological and biochemical findings are described and discussed in relation to available data from other pangolin species. These preliminary results provide the first reference framework for this species and highlight their relevance for clinical assessment, health monitoring, and conservation management.

## Introduction

Pangolins are the most trafficked mammals in the world, primarily due to the demand for their meat consumption and scales in traditional medicine (Xi al. 2025). Among the eight extant species, the giant pangolin (*Smutsia gigantea*) is the largest African pangolin species with a published record of a 42 kg male pangolin (Okala Research Team, 2022). This elusive species inhabits rainforests, swamp forests and wooded savannas across Central and West Africa (Hoffmann et al. 2020).

The giant pangolin is classified as Endangered on the IUCN Red List (IUCN, 2019) and faces severe population declines due to poaching and habitat loss. In countries of its range, local extinction has been projected within the two coming decades even in the absence of further habitat degradation (Zanva et al. 2023). Despite its conservation importance, the giant pangolin remains poorly studied, with major gaps in basic biological and clinical data. While conservation programs in Africa focus on pangolin rescue, rehabilitation and reintroduction, the absence of such data limits evidence-based veterinary management and disease monitoring.

Blood analysis, including hematological and biochemical parameters, is a fundamental tool in wildlife veterinary medicine, enabling health assessment, detection of pathological processes, and monitoring of physiological conditions (Didkowska et al. 2024). This is particularly important in species such as pangolins, which can conceal overt clinical signs until advanced stages of disease (Authors observations). Blood parameters provide insight into organ function, inflammatory processes, metabolic and nutritional status, and reproductive condition (Ding 2025). Although such data have been reported for some Asian and African species, they are absent for giant pangolins. Indeed, blood parameters references have been published for Sunda (Yindee et al. 2023; Yu et al. 2021; Ahmad et al. 2018) and Chinese (Khatri-Chhetri et al. 2015; Chin et al. 2015) pangolins. In contrast, for African species, hematological and biochemical information remains scarce. Reference data are available for Temminck’s pangolin (Connelly et al. 2020; Hooijberg et al. 2021) while only captive-based values have been described for the white-bellied pangolin (Kane et al. 2022).

Establishing baseline hematological and biochemical values for *Smutsia gigantea* is essential to support clinical assessments in rescue centers, enable early detection of health threats and physiological disorders, inform rehabilitation strategies, and strengthen long-term health monitoring of both wild and rehabilitated individuals. Here, we present the first published blood values for the giant pangolin, providing critical data for the conservation of one of the world’s most trafficked mammals.

## Materials and Methods

### Admission and Clinical management

On August 22, 2025, a giant male pangolin (later named “Akiba”) was confiscated from poachers in Campo Ma’an National Park (Cameroon). On August 26, 2025, he was transported 390 km via road transportation to the Tikki Hywood Foundation (THF) for rehabilitation and care.

Upon admission, the individual presented signs consistent with systemic compromise. The pangolin was found to be lethargic, unable to ambulate, and had reduced responsiveness to external stimuli with a notable absence of the defensive rolling response (Hoffmann et al. 2020). The individual’s inability to maintain a curled posture suggested a state of severe muscular weakness. Physical markers included poor body condition (emaciation) and muscle wasting, reduced skin turgor also suggested acute dehydration. Body weight at admission was 30.3 kg.

Akiba received supportive and intensive care, including assisted feeding and medical treatment. Enteral nutrition was provided once daily via tube feeding using a formulated diet, with progressive refeeding volume (from 120mL up to 600mL). Oral rehydration salts (100mL) were also administered during the first four days post-admission. Tube feeding procedure was performed under a light inhalation sedation plane (stage II) using Isoflurane (Piramal Critical Care, Telanga, India) at 5% dose delivered in medical oxygen (5 L min-1) via a face mask for 1 minute and 30 seconds. Pre-oxygenation for 5 minutes (1 L min-1) delivered in flow-by was performed prior to sedation to reduce any negative respiratory effects following isoflurane inhalation such as hypoventilation or apnea (van Oostrom et al. 2017).

Medical treatment included: a single dexamethasone injection (0.2mg/kg SC) for management of shock (Cicarelli et al. 2007, 2006), antimicrobial therapy with amoxicillin–clavulanic acid (8.75mg/kg SC, once daily for 5 days) to treat possible underlying bacterial infection regarding his compromised condition, and supplementation with thiamine and iron to prevent respectively refeeding syndrome (5mg IM, once daily for 5 days) (Mehanna et al. 2008) and iron-deficiency anemia (Jogu et Kamran 2026) as physical findings included fatigue and pallor (0.08mg/kg PO, once daily for 3 days). After one week of care, body weight increased to 31.4 kg and clinical condition significantly improved.

### Sampling procedure and Laboratory analyses

Blood sampling was performed on day 11 post-admission, when the animal was deemed clinically recovered and fit for release based on normal behavioral and clinical parameters: bright and alert demeanor, normal thoracic auscultation and respiratory pattern, pink mucous membranes, normal pace gait without imbalance, nocturnal activity, ability to raise body using forelimbs, effective digging behavior, and strong resistance to handling. Blood sampling was conducted as part of the pre-release medical assessment (Memorandum of understanding, 0022) to ensure the absence of major blood parameters abnormalities that could contraindicate release, based on comparisons with available data from other pangolin species.

Blood sampling was performed under light sedation during daily tube feeding procedure. Complete morphometric measurements were additionally performed according to parameters described by Perera et al. (2020) using a flexible measuring tape. Blood was collected in one EDTA tube and one dry tube by a veterinarian from the medial caudal venous complex using a 16-gauge needle and a 5 mL syringe. In addition, two blood smears were made and stained with a fast-acting variation of May-Grünwald Giemsa staining (RAL 555, Cella Vision). Blood samples were kept refrigerated for 12 hours before hematological and biochemical analyses were conducted at Prima Laboratoire (Yaoundé). Hematology was performed using an XN-330 analyzer (Sysmex, Switzerland), and serum biochemistry was analyzed with a Cobas Integra 400-Plus system (Roche, France).

### Data visualization

Akiba blood parameters were compared with blood available values for other pangolin species. As several datasets were available in Chinese, Sunda and Temminck’s pangolins, reference values used for interspecific comparison were selected with a hierarchical approach. Priority was first given to studies including clinically healthy individuals, as physiological status is a major driver of hematological and biochemical parameters. Among these, preference was then given to data obtained from free-ranging animals, as the data are more representative of natural conditions than rescued populations. Finally, if multiple studies met these criteria, those with larger sample sizes were prioritized. For Indian pangolins, the only available study involved a clinically compromised individual, this dataset was included for comparative purposes due to the absence of alternative reference values. Similarly for white-bellied pangolins, only a study based on captive individuals was available and included but comparisons should be interpreted cautiously. Reference intervals were preferentially used when available (Hooijberg et al. 2021; Yindee et al. 2023; Khatri-Chhetri et al. 2015). For studies lacking reference intervals, mean ± standard deviation values were reported (Kane et al. 2022). For single-case study, data were included without reference range (Mohapatra et al. 2014).

Comparative visualization of hematological and biochemical parameters across pangolin species was performed using R (v. 4.2.2). Plots were generated using the *ggplot2* package. For each parameter, mean values were represented using points and error bars indicating reference intervals or standard deviation depending on data availability. Amylase values were excluded from biochemical visualization due to scale differences that impaired graphical interpretation.

## Results

Hematological and biochemical findings in Akiba are presented in **Table 1**. Comparison of hematological and biochemical profiles across pangolin species are presented **Figure 1**. Akiba morphometric measurements are summarized in **Table 2**. Blood smear evaluation did not reveal any abnormalities (**Supplementary Figure 1**.).

**Table 1.**
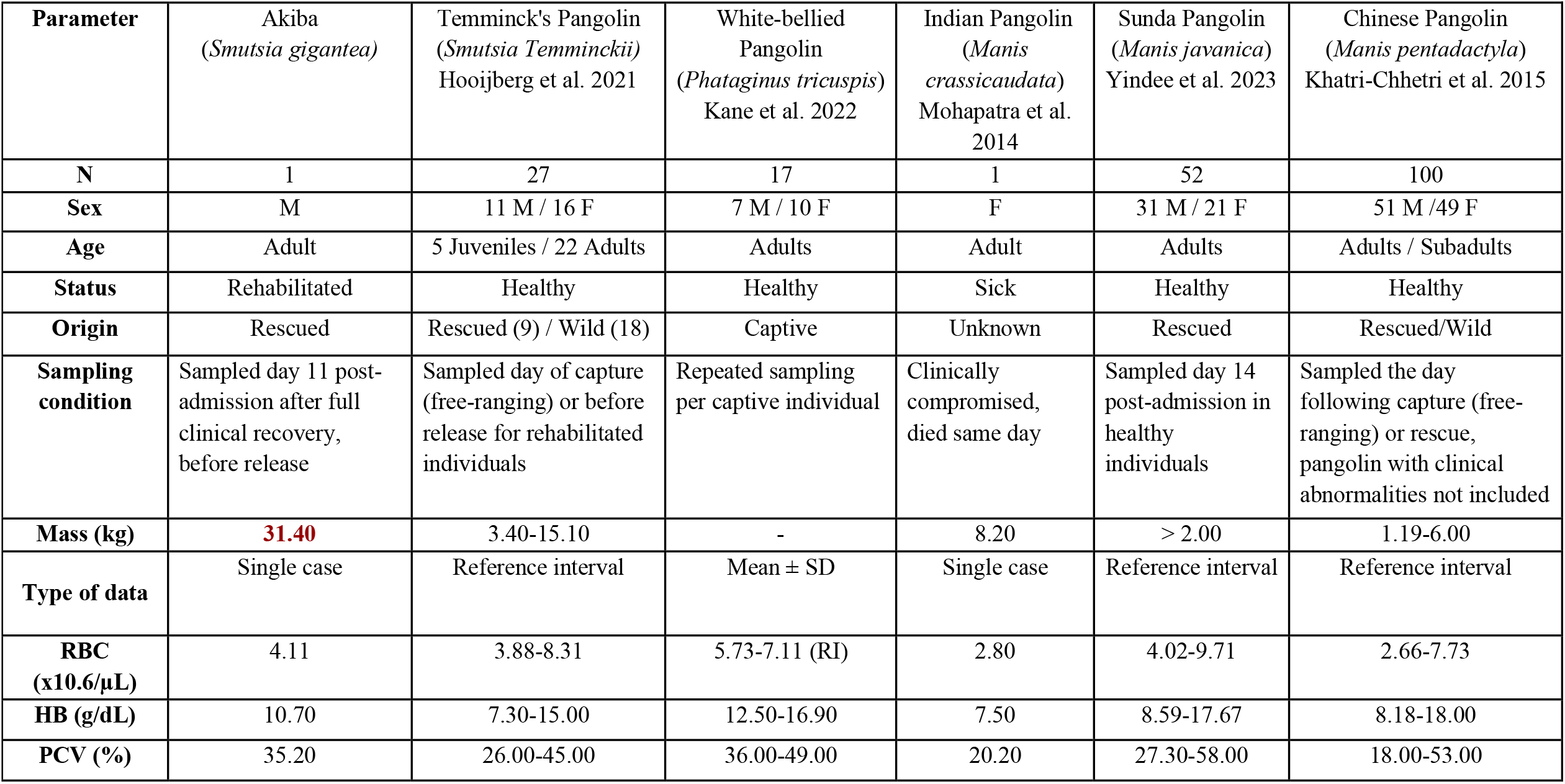
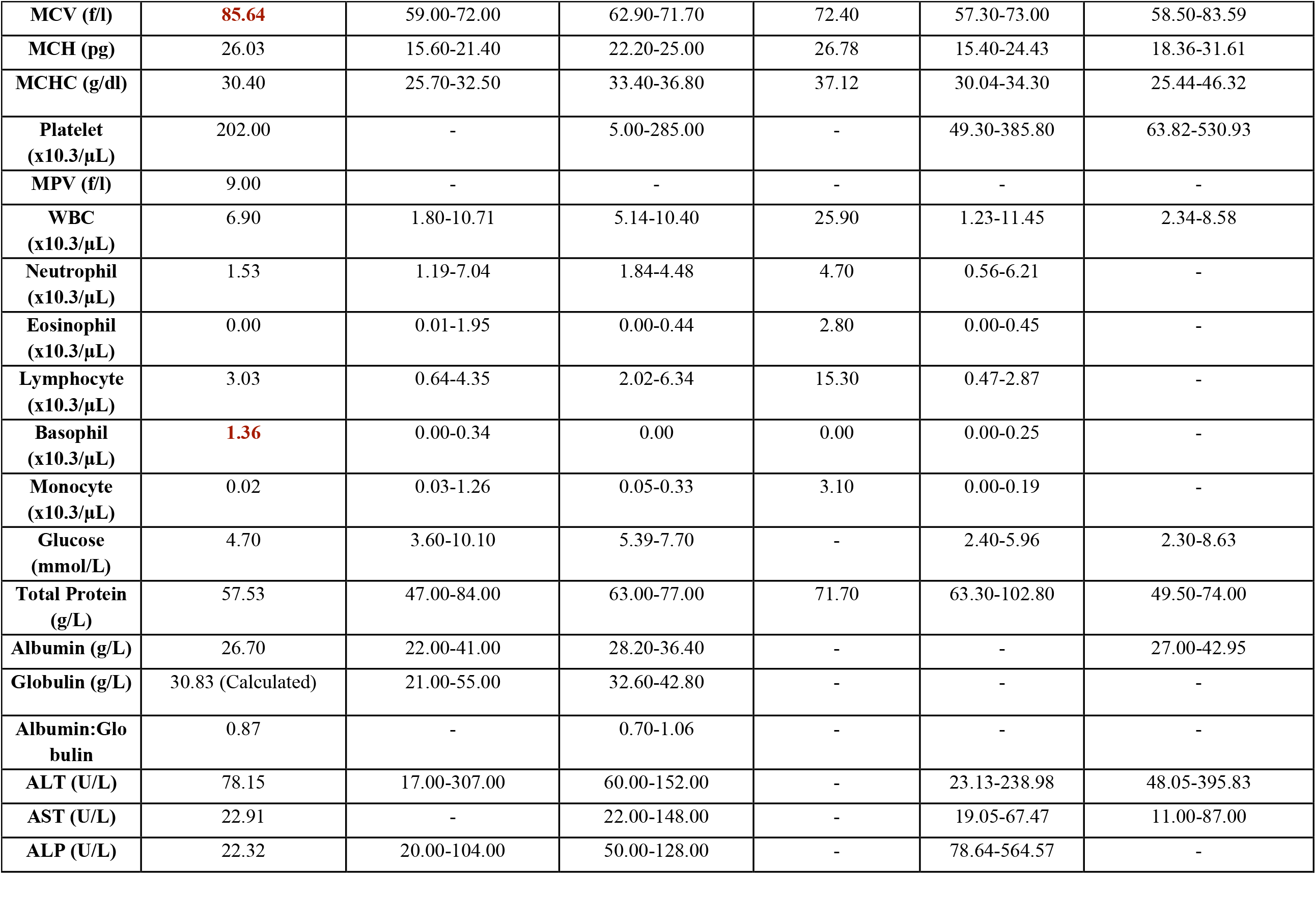
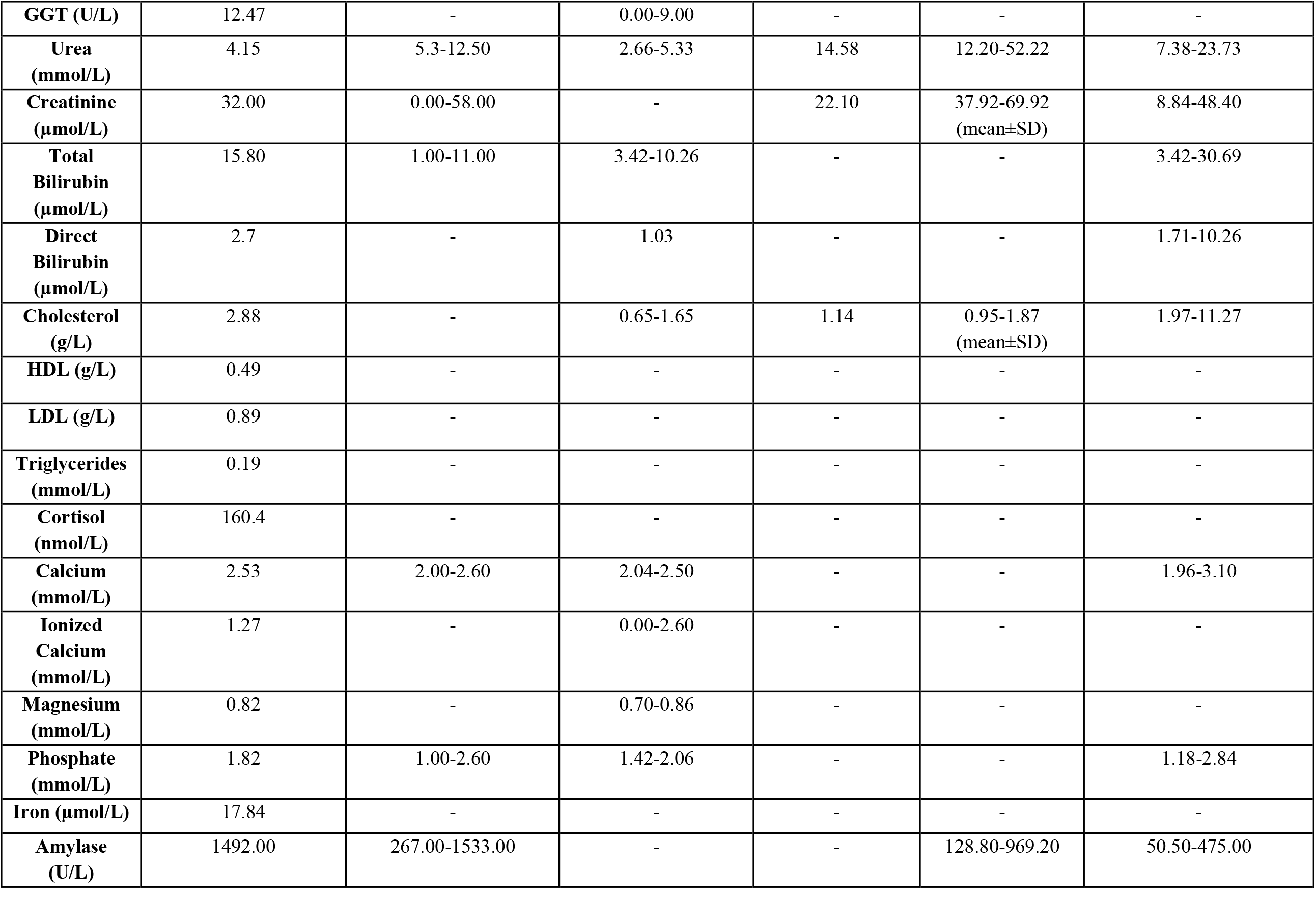
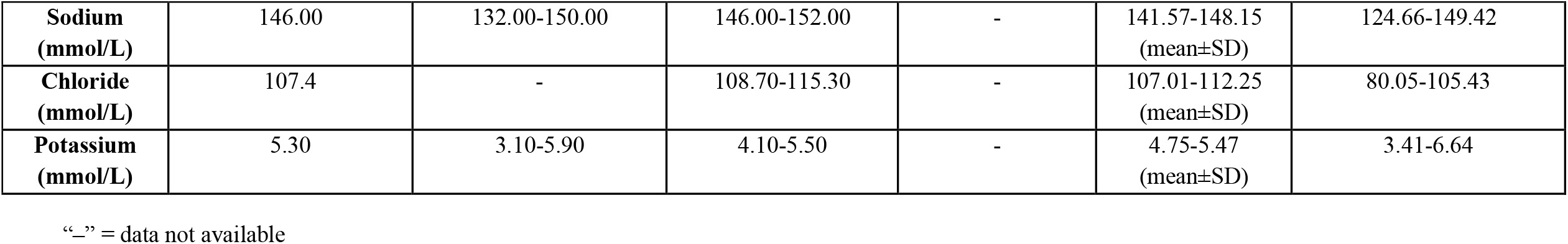
Haematologic and Biochemistry profile of “Akiba” compared to literature blood references across pangolin species. **RBC:** Red Blood Cell count. **HB:** Hemoglobin, **PCV:** Packed Cell Volume, **MCV:** Mean Corpuscular Volume, **MCH:** Mean Corpuscular Hemoglobin, **MCHC:** Mean Corpuscular Hemoglobin Concentration, **MPV:** Mean Platelet Volume, **WBC:** White Blood Cell count, **ALT:** Alanine amino-Transferase, **AST:** Aspartate amino-transferase, **ALP:** Alkaline Phosphatase, **GGT:** Gamma-Glutamyl Transferase. **Red:** parameter reported higher in giant pangolin compared to other previous species values.

**Figure 1.**
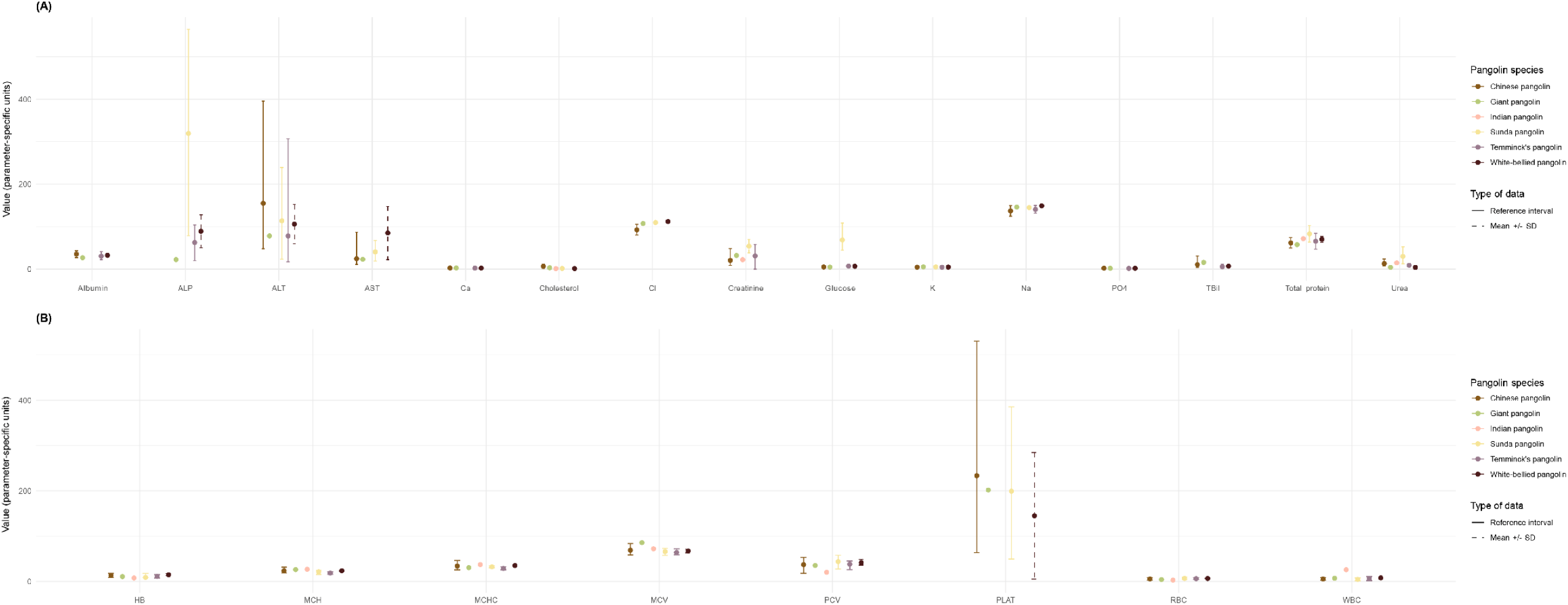
Comparison of biochemical **(A)** and hematological **(B)** parameters across pangolins species. Mean values were used as central estimates. When available, reference intervals are represented (solid lines); otherwise, mean ± standard deviation obtained from literature is displayed (dashed lines). Because reference intervals are not necessarily centered on the mean, error bars may appear asymmetrical. For species represented by a single individual, no error bars are shown.

**Table 2.** Morphometrics measurements of “Akiba” using Perera et al. (2020) morphometric data collection parameters. Head and body length was performed ventrally from tip of the nose to base of the anus. Additional measurements include total diameter when rolled into a ball, flesh part of the nose (from last scale to tip of the nose), ventral tail measurement (from base of the anus to tip of the tail), tail base girth and tail base width. Reported in cm. L: Left; R: Right.

|  |  |  |  |  |  |  |
| --- | --- | --- | --- | --- | --- | --- |
| Body Weight (BW) | 31.3 kg |  |  |  |  |  |
| Snout length (SL) | 9.7 |  |  |  |  |  |
| Flesh part of nose (FN) | 3.7 |  |  |  |  |  |
| Ear length (HL) | L: 3.1 |  |  | R: 3.2 |  |  |
| Head length (HL) | 21.4 |  |  |  |  |  |
| Neck circumference (NC) | 29.5 |  |  |  |  |  |
| Forelimb length (FLL) | 29.5 |  |  |  |  |  |
| Shoulder circumference (SC) | 77.0 |  |  |  |  |  |
| Body circumference (BC) | 83.0 |  |  |  |  |  |
| Head & body length (HBL) | Ventral: 65.0 |  |  |  |  |  |
| Hindlimb length (HLL) | 25.6 |  |  |  |  |  |
| Hind foot diameter (HFD) | L: 9.8 length / 7.6 width |  |  | R: 10 length / 7.4 width |  |  |
| Total Body Length (TBL) | 168.5 |  |  |  |  |  |
| Total Diameter in ball | 151.0 |  |  |  |  |  |
| Tail length (TL) | Ventral: 67.0 / Dorsal: 83.4 |  |  |  |  |  |
| Tail base girth (TG) | 51.0 |  |  |  |  |  |
| Tail base width (TW) | 19.0 |  |  |  |  |  |
| Forelimb claws curvature length (CL) | L | 2<sup>nd</sup>: 6.6 |  | 3<sup>rd</sup>: 9.6 | 4<sup>th</sup>: 7.5 |  |
|  | R | 2<sup>nd</sup>: 6.6 |  | 3<sup>rd</sup>: 9.4 | 4<sup>th</sup>: 7.4 |  |
| Hindlimb claws curvature length (CL) | L | 1<sup>st</sup>: 2.3 | 2<sup>nd</sup>: 3.8 | 3<sup>rd</sup>: 4.0 | 4<sup>th</sup>: 3.0 | 5<sup>th</sup>: 1.8 |

## Discussion

We report the first publication of hematological and biochemical data for endangered *Smutsia gigantea* from an adult male in Cameroon, undergoing rehabilitation at the Tikki Hywood Foundation. These results should be interpreted as preliminary descriptive values for the species.

The lack of blood data from giant pangolins likely reflects both ecological and logistical constraints. The species display limited distribution in African tropical forest with peculiar nocturnal and solitary ecology (Hoffmann et al. 2020), restraining field-based research. In addition, accessing the medial caudal vein in a clinically healthy individual without sedation is technically demanding due to the defensive behavior to roll into a ball, combined with the challenges associated with restraining and manipulating a large-size animal. In remote field conditions, maintaining appropriate sample preservation without refrigeration can also compromise biochemical integrity (Wu et al. 2017). Moreover, access to diagnostic laboratories equipped and willing to process wildlife samples remains limited in range countries. These constraints may explain the current gap in the literature but also highlight the value of this first documented case.

The number of biochemical parameters evaluated in our study exceed those reported in previous pangolin studies, notably including an extended electrolyte panel (Magnesium, Calcium, Phosphate, Iron), liver function markers (AST, ALT, ALP, GGT), as well as cholesterol and cortisol measurement which provide insights into metabolic, hepatic, and physiological status. Blood gas analysis and lactates measurement were not performed due to the inability to conduct immediate measurements under field conditions. Additional parameters described from other pangolin species such as creatine kinase (CK), and lipase could also have provided further information on muscle activity, and pancreatic function. While CK is a highly conserved enzyme that can be measured across species (Ferreira 2016; Ventura-Clapier et al. 1998), lipase results are more method-dependent and usually require species-specific immunological assays (Lim et al. 2022), which may lack reliability in non-model species such as pangolins.

Most of the parameters evaluated in our giant pangolin individual are included in previous reference intervals described in other pangolin species. Akiba’s hematological profile of packed cell volume (35.2%), red blood cell (RBC) count (4.11 × 10^6^/µL) and hemoglobin (10.70 g/dL) fall within the range reported for Temminck’s, Sunda and Chinese pangolins. However, Akiba’s RBC indices were characterized by increased mean corpuscular volume (MCV) and high range mean corpuscular hemoglobin (MCH) associated with low range mean corpuscular hemoglobin concentration (MCHC). These findings may reflect a regenerative response following physiological stress or nutritional imbalance during rehabilitation. Indeed, this pattern may occur in regenerative anemia, in which circulating immature erythrocytes (reticulocytes) are larger and contain proportionally less concentrated hemoglobin (Tvedten 2022). Elevated MCV is also commonly associated with deficiencies in cobalamin (vitamin B12) or folate (Killeen et Adil 2025). Alternatively, Akiba’s hematological profile may indicate species-specific differences related to body size. Indeed, giant pangolin is substantially larger than other African and Asian pangolin species, and interspecific variation in erythrocyte size has been documented across mammals, sometimes correlating with body mass and metabolic adaptations (Kozłowski et al. 2010).

The leucocyte profile of Akiba revealed a predominance of lymphocytes (Lymphocyte/Neutrophil ratio of 0.66). The documented Neutrophil-to-Lymphocyte Ratio (NLR) of pangolins did not reveal a clear pattern when compared across species, with marked variability observed. A lymphocyte dominant ratio was described in ex-situ captive white-bellied pangolins (Kane et al., 2022) and in a sick Indian pangolin (Mohapatra et al., 2014), whilst a neutrophil dominant ratio was described with higher upper reference value in Temminck’s, Chinese and Sunda pangolin species. NLR is widely used in clinical practice as a biomarker of systemic inflammation, elevated NLR is associated with increased inflammatory responses and various health conditions (Zahorec 2021). The extreme outlier leukogram reported in the Indian pangolin likely reflects pathological changes rather than physiological variation. Similarly, the higher hematological range of white-bellied pangolin values should be compared with further study on free-ranging individuals to remove captivity bias limiting the interpretation. Akiba’s hematological profile may indicate a lymphocyte-dominant leukogram in giant pangolin species, potentially reflecting a species-specific immune pattern. In mammals, lymphocytes are primarily associated with adaptive immunity, whereas neutrophils mainly mediate innate immune responses (Alberts et al. 2002). Alternatively, this profile could reflect a transient lymphocytosis and/or neutropenia rather than a physiological baseline characteristic.

Serum biochemical results revealed lower range urea and alkaline phosphatase (ALP) values compared to other pangolin species and may be explained by recent nutritional history, including fasting prior to rescue and subsequent artificial refeeding. Although these results are not necessarily pathological (Lum 1995; Salazar 2014), they emphasize the need for species-specific parameter references to enable accurate clinical interpretation in the giant pangolin.

Low urea concentration is commonly associated with reduced protein intake (Sun et al. 2016), while decreased ALP activity may also be linked to nutritional factors, including protein deficiency or inadequate intake of micronutrients such as Zinc and Magnesium (Ray et al. 2010). Experimental studies in mammals further indicate that dietary composition can influence ALP activity, notably vitamin D restriction or high-fat diets are associated with reduced enzyme activity (Oku et al. 2023).

Achieving adequate protein intake during rehabilitation remains particularly challenging in myrmeco-termitophagous species (Steinecker-Quast et al. 2024).

In addition, species-specific ecological and physiological traits may contribute to these findings. Giant pangolin has been reported feeding on larger ant and termite (>10 mm in length) species compared to other African pangolin (Difouo Fopa et al. 2020), potentially influencing digestive physiology and associated ALP activity. For instance, larger termites such as Macrotermes have high levels of proteins and lipids (Cheseto et al. 2024), and ALP plays a role in intestinal absorption notably by regulating lipid absorption, calcium intake, and maintaining intestinal barrier function (Santos et al. 2022). Moreover, gastrointestinal transit time generally increases with body mass across mammals (Karasov et al. 1986), suggesting possible slower digestive kinetics in the giant pangolin, which may contribute to distinct species-specific enzymatic activity patterns.

Conversely, amylase activity was in higher range values for Akiba than reported in other species. Amylase activity is typically elevated in animals with carbohydrate-rich diets which contrast with protein- and lipid-based diets of myrmeco-termitophagous mammals (Cheseto et al. 2024; Ayieko et al. 2012). In our case, increased amylase may reflect nutritional supplementation during rehabilitation notably the use of artificial diet. This hypothesis is further reinforced by previous observations of higher amylase activity in rehabilitated compared to free-ranging Temminck’s pangolin (Hooijberg et al. 2021), suggesting an effect of rehabilitation on this parameter. Less specifically, high amylase could reflect pancreatic, intestinal, or salivary gland disturbances, including inflammation or obstruction (Slack et al. 2010; Ben-Horin et al. 2002). However, no clinical abnormalities were observed in this individual. Without established species-specific reference intervals and upper limit of normal amylase, it remains premature to conclude hyperamylasemia in Akiba.

A marked basophilia was observed in the automated hematological analysis but was not confirmed on blood smear evaluation, highlighting potential analytical artefacts. Basophilia is classically associated with hypersensitivity reactions or parasitic infections (Sticco et al. 2026). In our case, no ectoparasites have been identified but stool flotation examination revealed the presence of strongyle-type eggs and coccidia, supporting exposure to gastrointestinal parasites (**Supplementary Figure 2**.). However, the difference between automated counts and smear interpretation suggests caution, as leukocyte differentials may be unreliable when using non-validated analyzers in non-model species. Morphological confirmation remains essential to avoid misinterpretation.

Comparisons of our data with other pangolin species must be interpreted with caution due to differences in study design, sample size, physiological status, and origin of individuals. Additionally, heterogeneity in data reporting (reference intervals, mean ± SD, or single-case values) further limits direct comparisons. Interpretation of the present hematological and biochemical findings must also account for pre-analytical and analytical sources of variation. Numerous animal factors including: age, sex, body mass, circadian variation and season, fasting status, medication and anesthesia, physical exercise and concurrent infection may influence blood parameters (Humann-Ziehank et Ganter 2012). Similarly, technique-related factors including sampling technique, tube selection, storage conditions, transport time, laboratory preparation and possible contamination before analysis at the laboratory may affect results prior to analysis (Humann-Ziehank et Ganter 2012). Importantly, analyses were performed using human-calibrated automated analyzers not validated for pangolins as to date, there are no previous validations, likely due to material limitation (Hooijberg et al. 2021).

This may introduce measurement bias, particularly for indices dependent on erythrocyte (e.g., MCV, MCH) and leucocytes morphology (Grebert et al. 2021); and for enzyme measurements sensitive to species-specific structural variation. Indeed, differences in isoenzyme structure, co-factor affinity and protein interactions in non-model species may lead to inaccurate activity measurements if species-specific validation is not performed (O’Brien et al. 2024). Until proper validation studies are conducted, these data should therefore be interpreted cautiously.

The main limitation of this study is the reliance on a single individual which limits interpretation of the data and should therefore be compared with results from further extensive studies. However we believe the value of data presented in this manuscript may aid future veterinary treatment of this species. Differences observed between species may reflect genuine ecological and physiological adaptations, but could also arise from nutritional and rehabilitation context, or analytical bias. Broader sampling across individuals, sexes, seasons, and physiological states will be necessary to establish robust reference intervals.

With the addition of the present report, baseline blood data is now available for the giant pangolin, representing an important step toward a more comprehensive comparative clinical framework across Pholidota. Hematological and biochemical data are still lacking for two pangolin species: the black-bellied pangolin *(Phataginus tetradactyla)* and the philippine pangolin (*Manis culionensis*). These findings emphasize the importance of opportunistic data collection in rare and threatened species.

## Conclusion

- First publication of hematological and biochemical results in *Smutsia gigantea*. These results help to initiate a blood comparative framework for the genus *Smutsia* and other African pangolin species.
- Caution should be taken before extrapolation of the result as study is based on one single individual during post-rehabilitation phase, further studies should include larger sample size including both sexes and different classes of age.
- These data provide baseline data that will help clinical assessments of endangered giant pangolin and strengthen conservation medicines initiatives towards the species.

## Ethical considerations

The animal was handled as part of a wildlife rescue and rehabilitation program conducted by the Tikki Hywood Foundation. THF has been rehabilitating pangolins since 1994 and works in close cooperation with government officials and law enforcement. Procedures, housing and monitoring of Akiba were conducted under a Memorandum of understanding between THF and Cameroonian government relating to the support, conservation and protection of pangolin in Cameroon (0022). All procedures were performed with the aim of minimizing stress and ensuring animal welfare according to European ethical laws. Blood sampling was performed by a veterinarian as part of standard procedures to aid the rehabilitation of the individual.

## Conflict of Interest

*The authors declare that the research was conducted in the absence of any commercial or financial relationships that could be construed as a potential conflict of interest*.

## Author Contributions

**MM** conducted data collection, data analyses, and wrote the manuscript; **JHC** contributed to manuscript writing and editing on the manuscript; **JV** conducted data collection; **MD, EC, LH** contributed to study design, funding acquisition, and editing of the manuscript.

## Funding

Blood analysis was performed thanks to financial and technical support of Ceva Wildlife Research Fund.

## Acknowledgments

Blood analysis was performed thanks to financial and technical support of Ceva Wildlife Research Fund, authors are thankful for their trust and partnership. The authors would like to thank Prima Laboratory (Yaoundé) for their collaboration in the investigation of pangolin’s blood parameters and their continuous support, notably General Director Dr. TANG Jean-Maximilien and Technical Director Dr. NDZE Raphael. Authors are also thankful to the Okala Research Team for their guidance and experience in the rescue and rehabilitation of Akiba: Dr. LEHMANN David and Ms. LEWIS-SMITH Ruth.

## Data Availability Statement

All the datasets generated for this study are presented in the main manuscript.

## Notes

### Competing Interest Statement

The authors have declared no competing interest.

