## Supplementary Figure 1 for "First hematological and biochemical data in a rehabilitated giant pangolin (*Smutsia gigantea)* from southern Cameroon"

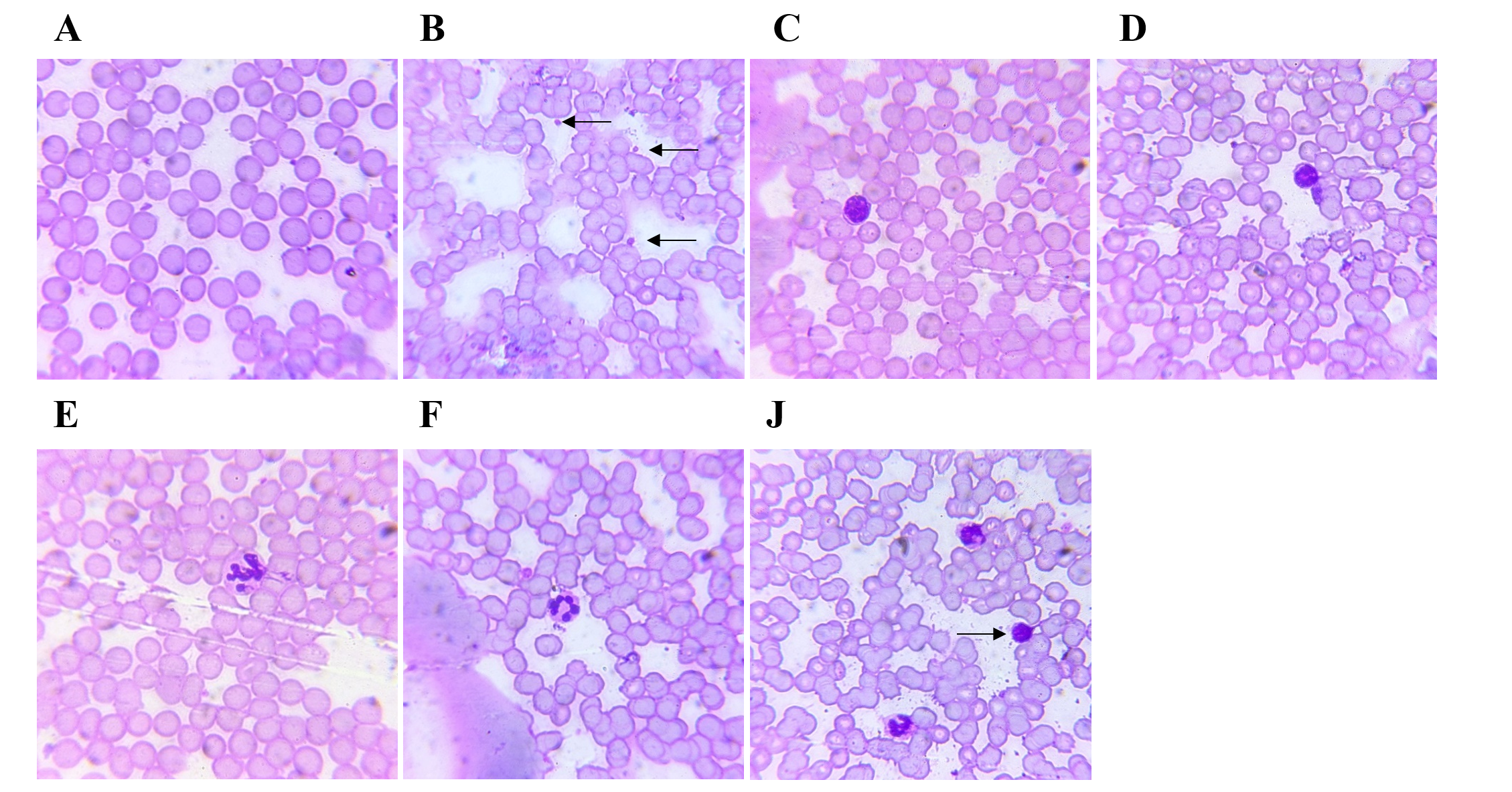

**Supplementary Figure 1.** Blood smear images from *Smutsia gigantea*. (A) Erythrocytes, (B) Platelets (arrow), (C) and (D) Lymphocytes, (E) and (F) Segmented Neutrophil, (G) Segmented neutrophils and Lymphocyte (arrow). May-Grünwald Giemsa, × 400 objective.
