## Supplementary Figure 2 for "First hematological and biochemical data in a rehabilitated giant pangolin (*Smutsia gigantea)* from southern Cameroon"

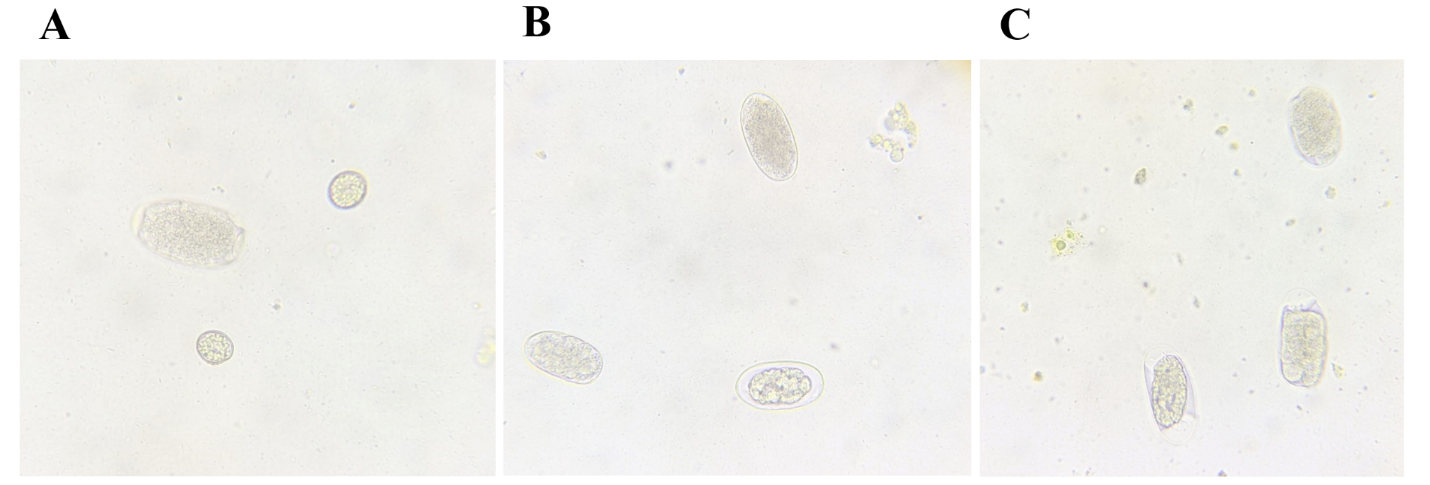

**Supplementary Figure 2.** Strongyle type eggs and coccidia identified during flotation examination of Akiba’s feces. (A) Strongyle type egg and two coccidia × 400 objective, (B) and (C) 3 strongyle type eggs × 400 objective.
